# Can natural selection favour indiscriminate spite?

**DOI:** 10.1101/745638

**Authors:** Matishalin Patel, Stuart A. West, Jay M. Biernaskie

## Abstract

Spiteful behaviours occur when an actor harms its own fitness to inflict harm on the fitness of others. Several papers have predicted that spite can be favoured in sufficiently small populations, even when the harming behaviour is directed indiscriminately at others. However, it is not clear that truly spiteful behaviour could be favoured without the harm being directed at a subset of social partners with relatively low genetic similarity to the actor (kin discrimination). Using mathematical models, we show that: (1) the evolution of spite requires kin discrimination; (2) previous models suggesting indiscriminate spite involve scenarios where the actor gains a direct feedback benefit from harming others, and so the harming is selfish rather than spiteful; (3) extreme selfishness can be favoured in small populations (and in some cases small groups) because this is where the feedback benefit of harming is greatest.

## Introduction

Spite is the hardest type of social trait to explain. Spiteful behaviour reduces the lifetime fitness of both the recipient and the performer (actor) of that behaviour (Hamilton 1970). In terms of Hamilton’s rule, –*C* + *RB* > 0, spite represents the case where there is a fitness cost to the actor (positive *C*), and a fitness cost to the harmed recipient (negative *B*), which can only be favoured if the genetic relatedness term, *R*, is negative. Understanding the meaning of negative relatedness is therefore crucial for explaining how and why spite evolves.

It has been argued that the evolution of spite requires kin discrimination, allowing the actor to direct harm towards a subset of individuals with whom they share relatively low genetic similarity (Foster & Ratineks 2000; Foster et al. 2001; Gardner & West 2004a,b, 2006; Gardner et al. 2004, 2007; Lehmann et al. 2006; West & Gardner 2010). Specifically, spite can be favoured when harming the less-similar individuals in a social group (primary recipients) reduces competition and therefore benefits the unharmed individuals (secondary recipients). In this case, negative relatedness arises because the actor’s genetic similarity to primary recipients is less than its genetic similarity to secondary recipients (Lehmann et al. 2006). In contrast, without kin discrimination, harming behaviours could not be directed at individuals to whom the actor is negatively related, so indiscriminate spite should be impossible.

However, a number of theoretical studies have suggested the possibility for indiscriminate spite. Hamilton (1970) originally suggested that if genetic similarity is measured relative to the entire population (including the actor), then there will be a negative relatedness between the actor and all others in the population, especially in small populations. Consequently, several papers have predicted that spiteful harming, directed indiscriminately at others, could be favoured in sufficiently small populations (Hamilton 1970, 1971; Grafen 1985; Vickery et al. 2003; Taylor 2010; Smead & Forber 2012). As a specific example, Verner (1977) and Knowlton and Parker (1979; Parker & Knowlton 1980) suggested that individuals could be favoured to hold territories that are larger than needed for their own interest (“super-territories”), in order to spitefully exclude others from resources. It is not clear, though, whether such indiscriminate harming traits are truly spiteful.

Here, we resolve this disagreement over indiscriminate spite. Many harming traits will be costly to primary recipients (*B* < 0) but provide a direct fitness benefit to the actor, because they reduce competition for the actor or its offspring. Consequently, the traits are selfish (–*C* > 0) rather than spiteful (–*C* < 0) (Hamilton 1970; Keller et al. 1994; Foster et al. 2001; West & Gardner 2010). We address the possibility that indiscriminate harming traits like territory size have been misclassified as spiteful when they are actually selfish (Colgan 1979; Tullock 1979). Our specific aims are to: (1) determine generally whether indiscriminate harming evolves as a spiteful or a selfish trait; (2) examine how different modelling approaches can change the meaning of negative relatedness and lead to misclassification of harming traits; (3) re-analyse the Knowlton & Parker (1979) territory-size model to determine whether it predicts spiteful behaviour.

### Harming traits

We first modelled natural selection acting on a harming trait, following the approach of Lehmann et al. (2006). The trait has a fitness effect on a focal actor (–*C*) and on two categories of recipients: the harmed primary recipients and the unharmed secondary recipients who benefit from reduced competition (fitness effects *B*_1_ and *B*_2_, respectively). Crucially, we define an individual’s fitness as its number of offspring that survive to adulthood (not simply the number of offspring produced), which is consistent with other definitions used for classifying social traits (Hamilton 1964; Rousset 2004; Lehmann et al. 2006; West et al. 2007). We assume that fitness effects on the actor, primary recipients, and secondary recipients must sum to zero because of competition for finite resources (Rousset & Billiard 2000):

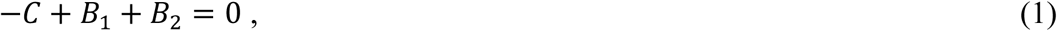

implying that any decrease in fitness for one category necessarily means an increase in fitness for another. Our model could apply to any finite population of constant size or to a local “economic neighborhood” (Queller 1994) in which there is a zero-sum competition for access to the next generation. Key examples of such local competition include polyembryonic wasps competing for resources inside a host (Gardner & West 2004a; Gardner et al. 2007) or male fig wasps competing for females inside a fig (West et al. 2001).

To predict the direction of natural selection acting on the harming trait, we considered the fate of a mutant harming allele in a population of individuals with a fixed, resident genotype. The success of the mutant allele depends on its “inclusive fitness effect” (Hamilton 1964): the sum of effects from a focal actor’s mutant trait on its own fitness and on the total fitness of each recipient category, weighted by their genetic similarity with the actor. Under the usual assumptions of weak selection and additive gene action, the inclusive fitness effect for our model is

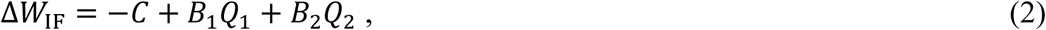

where *Q*_1_ and *Q*_2_ are probabilities of sharing identical genes between the focal actor and a random individual from the primary and secondary recipients, respectively. We note that the fitness effects in Equation 2 could alternatively be weighted by relatedness coefficients, where genetic similarity is measured with respect to a reference population (e.g., 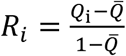, where 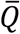 is the average genetic similarity in the entire population, including the actor; Hamilton 1970). However, doing this would not change any of the results given below. We therefore prefer the simpler approach used in Equation 2 and what follows below.

In the following sections, we examine two different ways of defining the category of secondary recipients and therefore partitioning the fitness effects of harming. Both methods correctly predict the direction of selection (they give the same sum as in Eq. 2). The first partitioning also maintains complete separation of direct and indirect fitness effects (–*C* and *RB*, respectively), making it appropriate for classifying harming traits as selfish (–*C* > 0) or spiteful (–*C* < 0). In contrast, the second partitioning obscures the separation of direct and indirect fitness effects, making it inappropriate for classifying traits in this way.

### Is indiscriminate harming spiteful or selfish?

We determined the conditions for a harming trait to be classified as spiteful or selfish. For this purpose, we assume that the focal actor, primary recipients, and secondary recipients are mutually exclusive categories. This ensures that the actor is not a recipient of its own behaviour, and so the –*C* term in the inclusive fitness effect (Eq. 2) captures all effects of the actor’s harming behaviour on its own fitness. From Equation 2, we derived the typical two-party version of Hamilton’s rule by eliminating the fitness effect on secondary recipients, using *B*_2_ = *C* – B_1_ (from Eq. 1). After rearrangement, the inclusive fitness effect is positive, and the harming trait is favoured, when

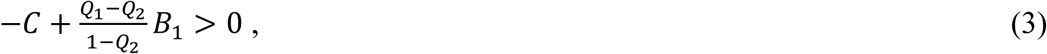

which is Hamilton’s rule with the relatedness between actor and primary recipients given by 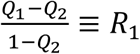. This is the genetic similarity between the actor and an individual from the potential primary recipients, measured relative to an individual from the potential secondary recipients.

Equation 3 implies that indiscriminate spite cannot evolve. This is because negative relatedness (and hence an indirect fitness benefit of harming) will arise only if harm can be directed at primary recipients who are less genetically similar to the actor than secondary recipients are (*Q*_1_ < *Q*_2_). In contrast, if the actor were harming others indiscriminately—for example, harming a random subset of a population or local economic neighbourhood—then its expected similarity to these primary recipients would be the same as to the set of potential secondary recipients (*Q*_1_ = *Q*_2_), and relatedness would be zero (*R*_1_ = 0). This implies that indiscriminate harming will be favoured when it is a selfish trait with a positive direct fitness benefit (–*C* > 0).

### Why does misclassification occur?

Misclassification of harming traits can occur because the fitness effects of social traits can be partitioned in different ways (Frank 1998). An alternative way of partitioning the effects of harming is to include the actor in the set of secondary recipients who may benefit from reduced competition. In fact, it is often implicitly assumed that the set of potential secondary recipients is the entire population (or economic neighbourhood), including the focal actor (Hamilton 1970, 1971; Grafen 1985; Vickery et al. 2003; Taylor 2010; Smead & Forber 2012). To make this explicit, we re-write the inclusive fitness effect as

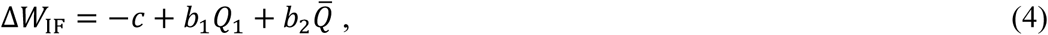

using lower-case letters to indicate that the fitness effects no longer match those from Equation 2. In particular, *b*_2_ is now the benefit of reduced competition that may be experienced by all individuals in population (including the actor), and 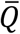 is the probability of genetic identity between the focal actor and a random individual the entire population (including itself). It follows that –*c* is not a total direct fitness effect because it excludes the secondary benefit of harming that feeds back to the focal actor (increased direct fitness due to reduced competition; Fig. 1).

**Figure 1.**
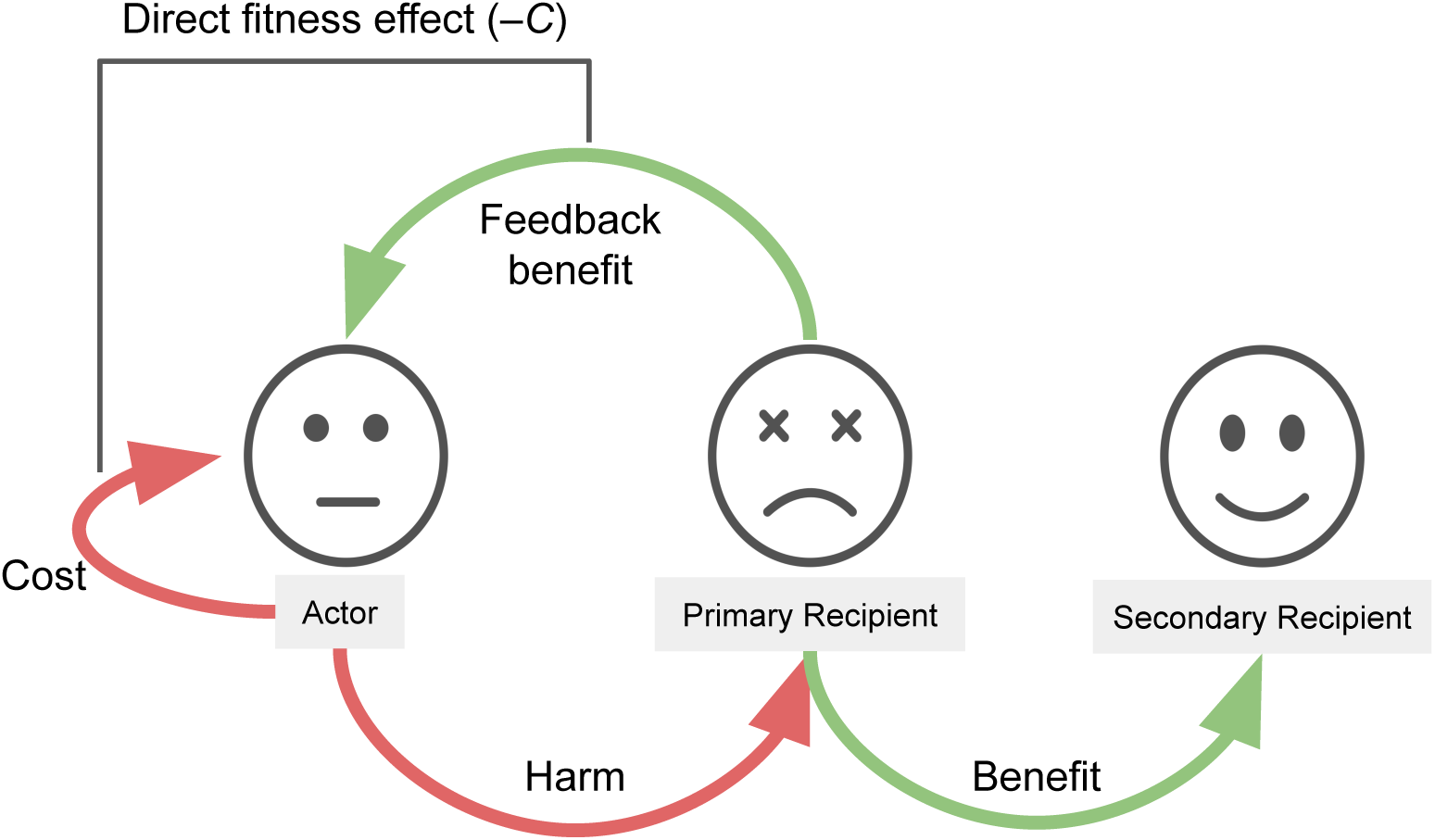
Partitioning the fitness effects of a harming trait. When a focal actor harms a primary recipient, this reduces competition and may therefore benefit the unharmed secondary recipients and the actor itself (“feedback benefit”). Some modelling approaches include the actor in the set of secondary recipients of the harming trait. However, the total direct fitness effect (–*C* in Hamilton’s rule) includes the fecundity cost of expressing the harming trait plus the feedback benefit.

We used Equation 4 to derive an analogue of Hamilton’s rule, which reveals a different version of negative relatedness. For example, in a population (or economic neighbourhood) of *N* individuals, an actor could indiscriminately harm a random subset of individuals with genetic similarity *Q*_1_ to the actor. If the entire population is in the set of secondary recipients, then the expected genetic similarity between the actor and these recipients is 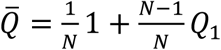 (where the first term accounts for the actor’s similarity to itself). Eliminating the fitness effect on secondary recipients (using *b*_2_ = *c* – *b*_1_), shows that indiscriminate harming is favoured when

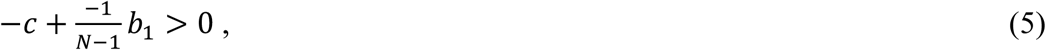

where –1/(*N* – 1) is the relatedness between actor and primary recipients, measured with respect to the entire population 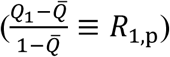. This is the version of negative relatedness that has led to predictions of indiscriminate spite in small populations (e.g., Hamilton 1971; Grafen 1985).

However, although the term 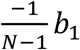 resembles an indirect fitness benefit (*RB* > 0), it actually accounts for the secondary fitness benefit of harming that feeds back to the focal actor. This can be made more explicit by deriving an analogue of Hamilton’s rule from Equation 4, this time eliminating the fitness effect on primary recipients (using *b*_1_ = *c* – *b*_2_). For example, in a well-mixed population of *N* individuals, indiscriminate harming is favoured when

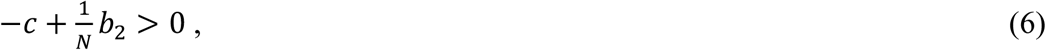

where 1/*N* is the relatedness between the actor and the entire population (including itself), measured with respect to primary recipients 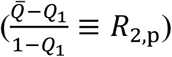. The term (1/*N*)*b*_2_ accounts for the fraction of the secondary benefit (reduced competition) that feeds back to the focal actor, which gets larger as the actor makes up a larger fraction of the population.

Our key distinction here is that harming behaviours can be either beneficial or costly to the actor (–*C* > 0 or –*C* < 0), whereas spiteful behaviours are strictly costly to the actor (–*C* < 0). We showed that indiscriminate harming is always favoured because it is beneficial to the actor— it has a positive effect on the actor’s number of surviving offspring (–*C* > 0). Moreover, indiscriminate harming can be favoured most in small populations (or small economic neighbourhoods) because this is where the focal actor can benefit most from the reduced competition that results from its harming behaviour.

### Re-visiting “super-territories”

We next re-examined the territory size model from Knowlton & Parker (1979; Parker & Knowlton 1980). We first analysed the model to fully separate direct and indirect fitness effects (applying Eq. 2), asking whether the model predicts selfish behaviour, as expected. We then used the alternative approach (applying Eq. 4) to illustrate why previous studies have interpreted territory size as a spiteful trait.

We considered a finite, deme-structured population (“island model”; Wright 1943) with *d* demes (assuming *d* > 1) and *n* individuals competing for territory in each deme (total population size is *N* = *dn*). Individuals that secure a territory have offspring and then die before a fraction *m* of their offspring disperse independently to a random deme in the entire population. All individuals have a genetically-determined strategy for the size of territory that they try to obtain (a continuous trait). Taking over a larger territory has three key effects: (1) it incurs a fecundity cost for the actor (we assume a linear cost with increasing trait size, with slope –*a* and *a* ∈ [0,1]); (2) it harms the actor’s deme mates by taking resources away and reducing their fecundity; (3) it reduces the competition faced by all remaining offspring in the population to secure a territory in the next generation.

We first assumed that the actor, primary recipients, and secondary recipients are mutually exclusive categories (as in Eq. 2). In the Appendix, we derive an expression for the fitness, *W*, of a focal actor as a function of its own territory-size strategy, *x*; the average strategy of its deme mates (primary recipients), *y*; and the average strategy of individuals in other demes (secondary recipients), *z*. We used this “neighbour-modulated” fitness function to derive the inclusive fitness effect, by taking partial derivatives with respect to the strategies of the different categories of individuals (Taylor & Frank 1996; Rousset & Billiard 2000):

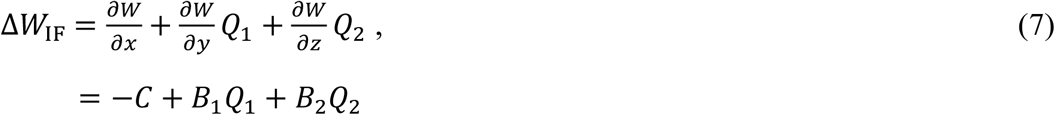

where all partial derivatives are evaluated in a monomorphic population (*x* = *y* = *z*). We derive expressions for *Q*_1_ and *Q*_2_ in the Appendix, and with these we determined the equilibrium of the model (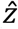, where directional selection stops) by solving Δ*W*_IF_ = 0. We also checked that the equilibrium is a convergence-stable strategy, denoted *z*\*, meaning that if the population is perturbed from the equilibrium then natural selection will push it back 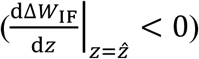.

We found that the equilibrium of our model, *z*\* = 1/(*aN*), is identical to that originally predicted by Parker & Knowlton (1980); however, our analysis shows that the optimal territory size strategy is selfish rather than spiteful. Territory size cannot be spiteful in this model because the actor’s genetic similarity to individuals in other demes is always equal to or less than the similarity to deme mates (*Q*_1_ ≥ *Q*_2_). Accordingly, the relatedness to primary recipients (measured relative to secondary recipients) is never negative (*R*_1_ ≥ 0), and so there is no indirect benefit of larger territory size. Moreover, when offspring dispersal is limited (*m* < 1) and deme mates are positively related (*R*_1_ > 0), there is no indirect benefit of smaller territory size (as a form of helping). This is because limited dispersal increases competition among offspring within the deme, which promotes harming and exactly cancels the effect of positive relatedness (as in Taylor 1992). Territory size therefore evolves for its direct benefit only, with larger territories promoted by a smaller fecundity cost to the actor (smaller *a*) and smaller population size (smaller *N*). Specifically, the direct fitness effect at equilibrium (*z* = *z*\*) is

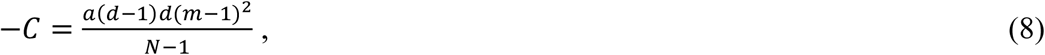

which is either positive (when *m* < 1) or zero (when *m* = 1). In the case of full offspring dispersal (*m* = 1), the equilibrium is the point where the fecundity cost to the actor is exactly balanced by the feedback benefit experienced by its offspring (reduced competition for space in the next generation). As the population approaches this equilibrium, however, direct fitness is always positive (–*C* > 0), confirming that territory size evolves as a selfish trait (Fig. 2).

**Figure 2.**
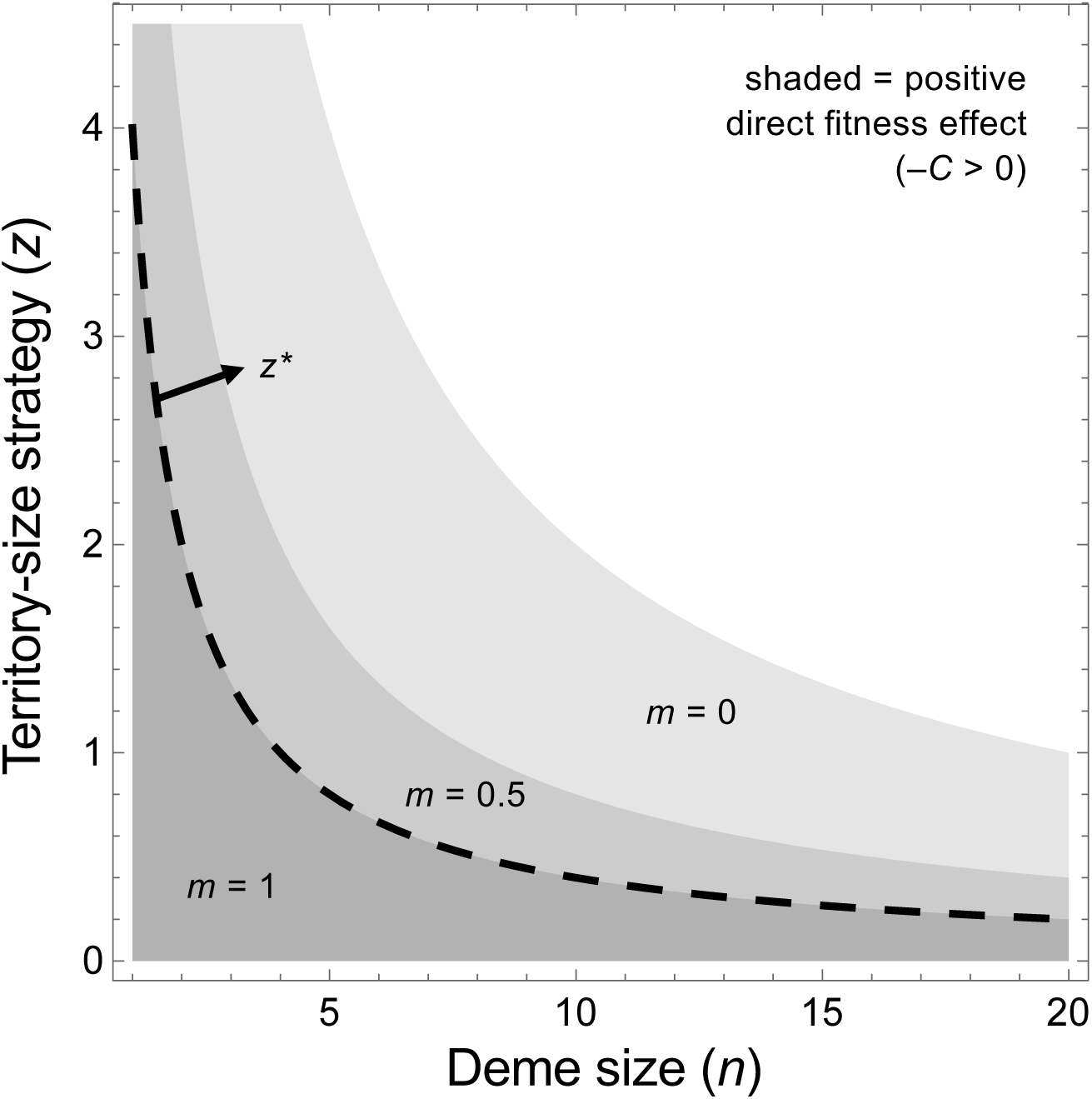
Territory size and direct fitness. Larger territory size is promoted by smaller population size (smaller *dn*) and reduced offspring migration from the deme (smaller *m*), both of which increase the direct benefit to an actor for harming its deme mates. However, reduced migration also increases the relatedness among deme mates, which inhibits larger territory size. Ultimately, the optimal territory size strategy (*z*\*, dashed line) is independent of migration rate and evolves as if the population were fully mixed (*m* = 1). Other parameters used: *d* = 5, *a* = 0.05.

We next assumed that the set of secondary recipients is the entire population, including the focal actor (as in Eq. 4). In this case, the inclusive fitness effect is

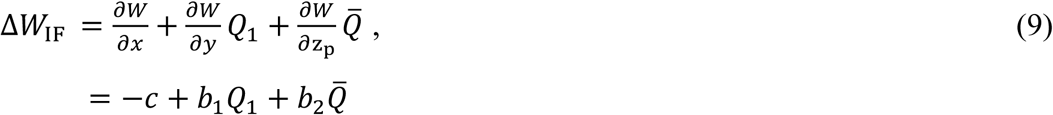

where *z*_p_ is the average territory size strategy in the entire population (including the focal actor), and all partial derivatives are evaluated at *x* = *y* = *z*_p_. As expected, solving for the equilibrium of Equation 9 gives the same answer as before, *z*\* = 1/(*aN*).

This version of the model shows, however, why territory size could be misclassified as spiteful. For example, in a fully mixing population at the equilibrium (*m* = 1; *z*_p_ = *z*\*), the first term in Equation 9 is

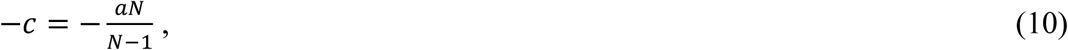

which is always negative. This term reflects the fecundity cost of the focal actor’s territory size strategy; however, it is not the total direct fitness effect because it excludes the feedback benefit experienced by the actor’s offspring (reduced competition). As noted above, when *m* =1 this feedback benefit should exactly balance the fecundity cost at equilibrium. Following Equations 5 or 6, we can calculate the feedback benefit as (–1/[*N*–1])*b*_1_ or (1/*N*)*b*_2_ (both evaluated at *z*_p_ = *z*\*), which gives the expected result, *aN*/(*N* – 1). The partitioning in Equation 9 therefore splits the total direct fitness effect of territory size into two separate terms, –*c* + (–1/[*N*–1])*b*_1_ or –*c* + (1/*N*)*b*_2_, which could be misinterpreted as a direct fitness cost (–*C* < 0) and an indirect fitness benefit (*RB* > 0).

## Discussion

We examined a general model of harming traits and a specific model where larger territory size is an indiscriminate harming trait. In both models we found that: (1) the evolution of spite requires kin discrimination, where the actor harms only a subset of other individuals (those with relatively low genetic similarity); (2) without kin discrimination, harming can be favoured but only when there is a sufficient direct, feedback benefit to the actor (reduced competition for the actor or its offspring); (3) indiscriminate harming can be favoured most in small populations (or small economic neighbourhoods), where the feedback benefit to the actor is greatest; (4) previous studies have misclassified indiscriminate harming as spite, partly because they misinterpret the feedback benefit as an indirect (kin-selected) benefit (*RB* > 0). Overall, our analyses illustrate why indiscriminate harming traits are selfish rather than spiteful.

### Classifying harming traits

For the purposes of classifying harming traits, we found that it is easiest to treat the actor, primary recipients, and secondary recipients as separate categories. This makes it straightforward to separate the total direct and indirect fitness effects of harming (–*C* and *RB*, respectively) and ensures that non-zero relatedness will always be associated with an indirect fitness effect. For example, spiteful harming (–*C* < 0, *B* < 0) requires that harm is directed at primary recipients to whom the actor is negatively related (with respect to secondary recipients; *Q*_1_ < *Q*_2_ and *R*_1_ < 0), resulting in a positive indirect fitness effect (*R*_1_*B* > 0) (Lehmann et al. 2006). In contrast, when harming is indiscriminate, the actor has zero relatedness to primary recipients (with respect to secondary recipients; *Q*_1_ *= Q*_2_ and *R*_1_ = 0), and so harming can be favoured as a selfish trait only (–*C* > 0, *B* < 0).

We showed that misclassification of indiscriminate harming is due to an implicit assumption that the focal actor is a secondary recipient of its own behaviour (Hamilton 1970, 1971; Grafen 1985; Vickery et al. 2003; Taylor 2010; Smead & Forber 2012). This means that some of the actor’s direct benefit of harming has been accounted for by a fraction of the fitness effects on recipients, giving the appearance of an indirect benefit (*RB* > 0). For example, in a well-mixed population where all individuals (including the actor) are considered secondary recipients, a fraction of the fitness effect on primary recipients (–1/[*N* – 1] *B*_1_) actually contributes to the direct benefit of indiscriminate harming.

Others have suggested that harming traits should be classified based on their primary effects only, rather than their total fitness effects (Krupp 2013). This means that indiscriminate harming traits like larger territory size, which may be associated with a survival or fecundity cost (–*c* < 0 in the terms of our model), would be classified as spiteful, despite the feedback benefit to the focal actor. We argue, however, that a classification based on total fitness effects (–*C* and *RB*) is more useful (Hamilton 1964; West et al. 2007). This is because it emphasises the fundamental distinction between spiteful harming, which is favoured by indirect fitness benefits and requires kin discrimination, versus selfish harming, which is favoured by direct fitness benefits and does not require kin discrimination (West & Gardner 2010). Similar arguments have been made for maintaining the distinction between altruistic helping (–*C* < 0, *B* > 0) and mutually-beneficial helping (–*C* > 0, *B* > 0) (West et al. 2007).

### Indiscriminate harming in nature

We found that selfish indiscriminate harming can be favoured most in small populations or small economic neighbourhoods (e.g., small groups with relatively local competition). This is because harming primary recipients leads to reduced competition for all individuals in the population or group, and a focal actor receives a larger fraction of this secondary benefit when it makes up a larger fraction of the population or group. Indiscriminate harming can therefore be thought of as producing a type of public good for secondary recipients (Tullock 1979), analogous to indiscriminate helping, which is often thought of as a public good for primary recipients. A key difference is that indiscriminate helping is inhibited by local competition (Taylor 1992; Griffin et al. 2004); in contrast, indiscriminate harming requires local competition so that the focal actor can actually benefit the reduced competition that results from its harming (Gardner & West 2004b).

So where can we expect to find the most extreme examples of selfish harming? As recognised by Hamilton (1970), very small populations will tend to extinction, so harming traits in these populations are unlikely to be observed. But examples of extreme selfishness should be found in small groups with relatively local competition, such that harming other individuals significantly reduces competition for the actor. One potential example is in fig wasps, where males fight for access to females, and the intensity of fighting increases sharply as the number of males in the fig declines (Murray et al. 1989; Reinhold 2003; West et al. 2001). Further potential examples include competition among female honey bees for a colony and other cases where males engage in local competition for mates (e.g., *Melittobia* parasitoids; Griffin & West 2002).

## Acknowledgements

We thank Guy Cooper, Asher Leeks, Alan Grafen, and Tom Scott for comments on the manuscript.

## Author contributions

All authors conceived and designed the study, MP drafted the initial version of the manuscript, and all authors contributed to later versions of the manuscript.

## Appendix: Territory-size model

### Deriving the fitness function

Here, we derive an expression for the fitness of a focal actor with a mutant territory size strategy, based on the models of Knowlton and Parker (1979; Parker and Knowlton 1980). We consider a population that is structured into *d* demes of *n* individuals competing for territories, where each deme has *A* units of available territory. The focal actor’s strategy, *x*, represents a continuous number of territory units that it attempts to gain (*x* > 0). The average strategy of the actor’s deme mates is *y*, and the average strategy in all other demes is *z*.

We first calculate the expected offspring production (expected fecundity, *F*) for the focal actor, an individual in the actor’s deme, and an individual in another deme. These expected values depend on: (1) the probability of an individual acquiring a territory (assuming that available spaces are acquired completely randomly); (2) the cost associated with the individual’s strategy (assuming fecundity declines linearly with increasing territory size strategy; *f* (*x*) = 1 – *ax*, where 0 < *a* < 1). For the focal actor, there are *A*/*y* spaces available in the deme, and we use the simplifying assumption that a mutant individual has priority to claim the territory units denoted by its strategy (Knowlton and Parker 1979). Therefore, the focal actor has a 1/*n* probability of acquiring a territory, and its expected fecundity is

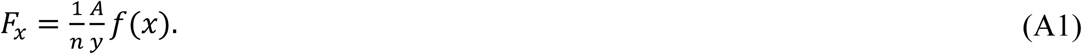

The space available for others in the patch depends on whether or not the focal actor claims a territory. The actor gains access to the patch with probability *A*/*ny*, and in this case (*A* – *x*)/*y* spaces remain; otherwise, *A*/*y* spaces are available. The expected fecundity for one of the *n* – 1 deme mates of the focal actor is therefore

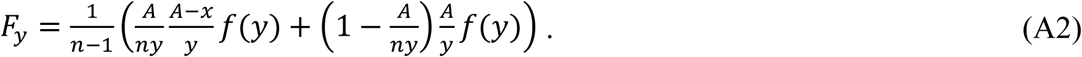

Finally, for an individual in another deme in the population, there are *A*/*z* spaces available, and so the expected fecundity for one of these individuals is

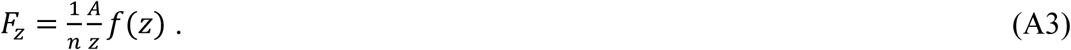

We next calculate the focal actor’s fitness, *W*(*x, y, z*), which is the number of its offspring that survive to compete for a territory in the next generation. This can be partitioned into two terms, the first term accounting for offspring that compete on the focal actor’s natal deme (those that did not disperse, with probability 1–*m*, and those that dispersed but landed on the natal deme, with probability *m*/*d*) and the second term accounting for offspring that disperse with probability *m* to compete in the *d* – 1 non-natal demes:

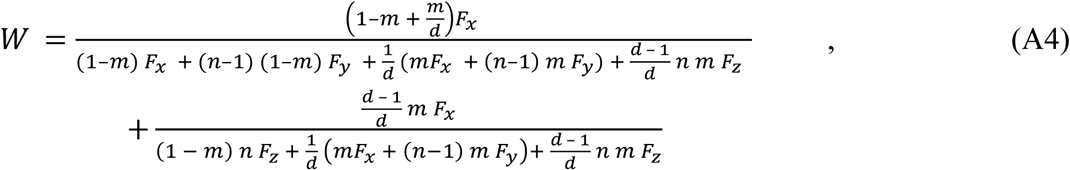

where the denominator of the first and second terms account for, respectively, all offspring competing in the focal actor’s natal deme and all offspring competing in any other deme in the population. Equation A4 is the fitness function used to calculate the inclusive fitness effect in Equation 7 of the main text. To express the focal individual’s fitness in terms of *x, y*, and *z*_p_ (the average territory size strategy in the entire population, including the focal individual), we substituted (*x* + (*n* –1)*y* – *dnz*_p_)/(*n* – *nd*) for *z* in Equation A4. This gives the fitness function used to calculate the inclusive fitness effect in Equation 9 of the main text.

### Deriving probabilities of genetic identity

Next, we derive probabilities of genetic identity by descent in a finite deme-structured population, following the approach of Taylor et al. (2000). In particular, we needed the probability of identity between the focal actor and a randomly selected deme mate (*Q*_1_), between the actor and a randomly selected individual in another deme (*Q*_2_), and between the actor and a randomly selected individual in the entire population (including itself), defined as

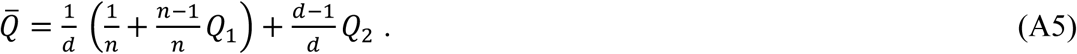

The remaining probabilities of identity are given by the following recursive equations:

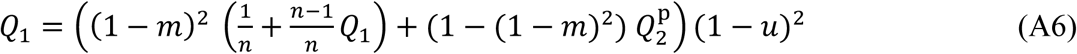

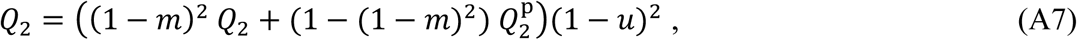

where *u* is the “contrived mutation rate” from Taylor et al. (2000). We solved Equations B1-B3 simultaneously and evaluated the solution in the limit of a low mutation rate (*u* → 0), giving:

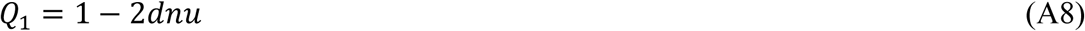

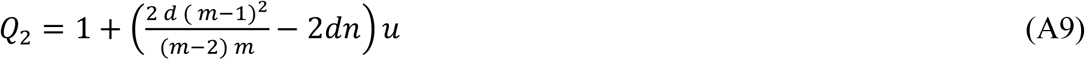

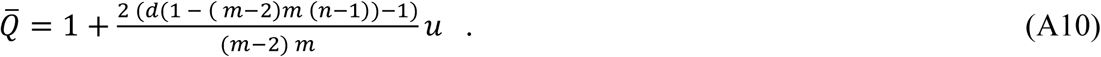

These are the probabilities of genetic identity used in Equations 7 and 9 of the main text.

